# Differential roles of CTP synthetases CTPS1 and CTPS2 in cell proliferation

**DOI:** 10.1101/2023.03.29.534741

**Authors:** Norbert Minet, Anne-Claire Boschat, Rebecca Lane, David Laughton, Philip Beer, Hélène Asnagli, Claire Soudais, Tim Bourne, Alain Fischer, Emmanuel Martin, Sylvain Latour

**Author notes:** **Corresponding author:** Sylvain Latour, Laboratory of Lymphocyte Activation and Susceptibility to EBV infection-Institut des Maladies Génétiques-Inserm UMR 1163, 24 boulevard du Montparnasse, 75015 Paris, France.

## Abstract

The CTP nucleotide is a key precursor of nucleic acids metabolism essential for DNA replication. De novo CTP production relies on CTP synthetases 1 and 2 (CTPS1 and 2) that catalyze the conversion of UTP into CTP. CTP synthetase activity is high in proliferating cells including cancer cells, however, the respective roles of CTPS1 and CTPS2 in cell proliferation are not known. By inactivation of CTPS1 and/or CTPS2 and complementation experiments, we showed that both CTPS1 and CTPS2 are differentially required for cell proliferation. CTPS1 was more efficient in promoting proliferation than CTPS2, in association with a higher intrinsic enzymatic activity that was more resistant to inhibition by 3-Deaza-uridine, an UTP analog. The contribution of CTPS2 to cell proliferation was modest when CTPS1 was expressed, but essential in absence of CTPS1. Public databases analysis of more than 1,000 inactivated cancer cell lines for CTPS1 or CTPS2 confirmed that cell growth is highly dependent of CTPS1 but less of CTPS2. Therefore, our results demonstrate that CTPS1 is the main contributor to cell proliferation.

## INTRODUCTION

As the building blocks of RNA and DNA, and as substrates of various cellular processes, nucleotides are key to most normal and pathological cellular and metabolic processes. Nucleotide levels are therefore tightly regulated, including through *de novo* synthesis, recycling and/or salvage pathways. The pyrimidine nucleotide cytidine triphosphate (CTP) is known to be in limiting concentrations in cells, in contrast to the other “main” nucleotides (ATP, GTP, TTP and UTP). CTP has also been linked to the synthesis of phospholipids and the sialylation of proteins [1–3]. Furthermore, it was recently shown that the limited availability of CTP shapes viral evolution [4] and CTP is a substrate for viperin, an enzyme that catalyzes the formation of ddhCTP, an antiviral nucleotide [5].

CTP arises from two sources, one dependent on the nucleotide salvage pathway, while the second involves *de novo* synthesis [6]. The salvage pathway recycles the nucleoside cytidine, a product of nucleic acid degradation. Cytidine is transformed into CTP via the synthesis of CMP and CDP. The de *novo* synthesis pathway of CTP is dependent on the enzymatic activity of CTP synthetases (CTPS). The CTPS activity is a two-step reaction involving two distinct enzymatic activities, a kinase and a glutaminase activity. CTPS catalyses the ATP-dependent amination of UTP into CTP using ammonia (NH_3_) transferred from the parallel hydrolysis of glutamine. In humans, CTPS activity is dependent on two enzymes with highly conserved structural identity, the CTP synthases 1 and 2 (CTPS1 and CTPS2) respectively encoded by the *CTPS1* and *CTPS2* genes. *CTPS1* and *CTPS2* share more than 75% of identity and are relatively well-conserved through species [7]. CTPS1 and CTPS2 contain two enzymatic domains, a synthetase/kinase domain and a glutaminase domain separated by a linker region. The C-terminal part is a regulatory domain with several sites of phosphorylation and is the most variable region between CTPS1 and CTPS2. CTPS1 is regulated by phosphorylation [8–10], ubiquitination [11] and polymerization [12,13]. CTPS1 forms dimers and tetramers representing inactive and active forms of the enzyme, respectively. Moreover, tetramers of CTPS1 can polymerize into higher order structures known as filaments, rod and rings or *cytoophidia*, whose function remains debated [14–17]. These different steps of CTPS1 containing supramolecular structure are influenced by the availability of substrates and products. Regarding the regulation of CTPS2, little is known. Two studies have reported that CTPS2 shares with CTPS1 at least some of these regulatory mechanisms, including phosphorylation and filament formation [18,19].

Both CTPS1 and CTPS2 proteins are expressed in all tissues (www.biogps.org; www.proteinatlas.org). CTP synthase activity is considered to be low in normal tissues while higher in proliferating tissues such as tumor cells, likely allowing malignant cells to overcome the CTP concentration bottleneck [20–22]. However, limited information is currently available on the respective role of CTPS1 and CTPS2 in proliferation. Notably, evidence of the roles of CTPS1/2 in proliferation was obtained by indirect approaches using inhibitors of CTPS activity [23,24]. Furthermore, these inhibitors have limited specificity with side effects on other metabolic pathways [25]. To date, the first and only direct evidence of the role of CTPS activity in cell proliferation has been provided by the recent identification of a cohort of immunodeficient patients harbouring a deleterious homozygous mutation in CTPS1 that results in a strongly reduced CTPS1 expression and activity [26,27]. CTPS1 expression was found to be upregulated in T lymphocytes following stimulation through antigen receptors, and necessary for their expansion during antigen-specific immune response. Importantly, CTPS1 was shown to be selectively required for the proliferation of activated T lymphocytes but not for their differentiation in effector cells.

To better understand the role respective of CTPS1 and CTPS2, we examined the impact of *CTPS1* and/or *CTPS2* gene inactivation by CRISPR-Cas9 on cell proliferation, viability and overall CTPS activity in two cell models and using recombinant proteins. We show that *CTPS1* and *CTPS2* are partially redundant and not equivalent to promote cell proliferation in correlation with differences in their enzymatic activity. Analysis of public databases of more than 1,000 inactivated cancer cell lines for *CTPS1* or *CTPS2* confirmed that cell growth is highly dependent on CTPS1, but not or less on CTPS2. In conclusion, our study documents that CTPS1 and CTPS2 are critical factors of cell proliferation with partially redundant roles.

## RESULTS

### CTPS1 and CTPS2 expression in different cell lines

In order to characterize the respective roles of CTPS1 and CTPS2 in cell proliferation, we first determined the levels of the CTPS1 and CTPS2 proteins in a variety of cell lines of hematopoietic origin as well as in the non-hematopoietic embryonic kidney cell line HEK-293T (hereafter designated as HEK) by western blot and RT-qPCR in a variety of cell lines of hematopoietic origin (**Fig. 1**). CTPS1 was found to be expressed at protein level in all tested cell lines, while CTPS2 expression was variable depending on cell origin. CTPS2 was not detectable or weakly expressed in some cancer cell lines of T lymphoid origin like MOLT-4 (a human T lymphoblast line from an acute lymphoblastic leukemia), HUT-78 (derived from cutaneous T lymphocytes from a patient with Sezary Syndrome) and Jurkat (an acute T-cell leukemia) cells, while it was detectable in CCRF-CEM cells, an acute lymphoblastic T-cell leukemia/T-ALL (**Fig. 1A**). CTPS2 was not detectable in the U937 cell line of myeloid origin. All B lymphoblastoid cell lines expressed CTPS2 as well as the NK92, K562 and THP-1 cell lines of NK, erythroid and myeloid origin, respectively. HEK cells expressed both CTPS1 and CTPS2. Expression of CTPS2 was higher in HEK than in cell lines of hematopoietic origin. Levels of *CTPS1* and *CTPS2* transcripts analysed by RT-qPCR in several of these cell lines were consistent with protein expression (**Fig. 1B**). Notably, HEK cells exhibited the highest level of *CTPS2* transcripts and ratio of *CTPS2*/*CTPS1* mRNA.

**FIGURE 1.**
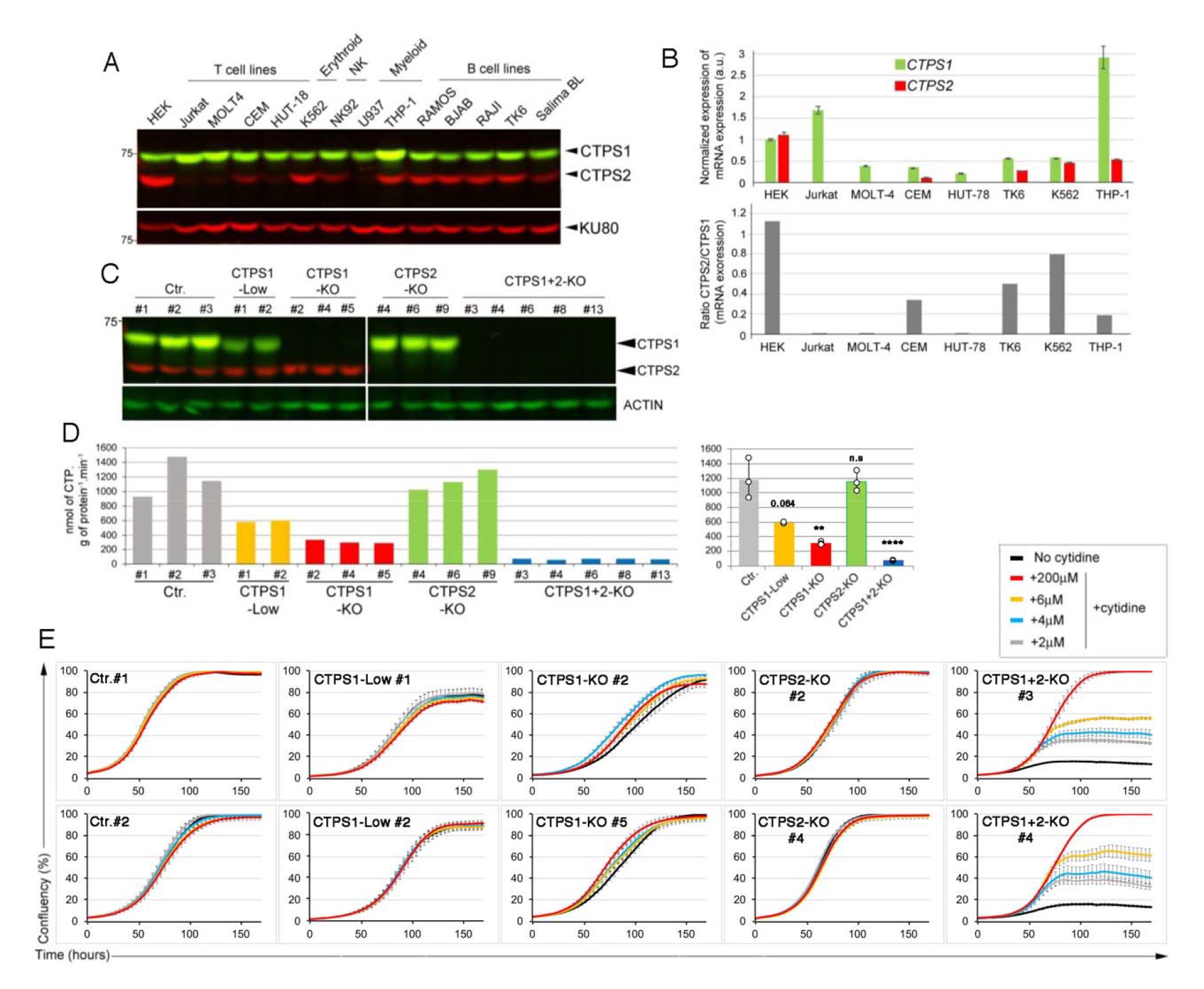
Differential roles of CTPS1 and CTPS2 in proliferation of HEK cells. **(A-B)** Expression of CTPS1 and CTPS2 in different cell lines. **(A)** Western blots of total cell lysates showing CTPS1 and CTPS2 expression in different hematopoietic cell lines with the exception of HEK that is a non-hematopoietic cell line. KU80 as the loading control. (**B**) Expression of *CTPS1* and *CTPS2* mRNA normalized to *GAPDH* mRNA by RT-qPCR (Upper panel). Ratio of *CTPS2*/*CTPS1* mRNA expression from RT-qPCR data (Lower panel). (**C**) Western blots showing the expression of CTPS1 and CTPS2 in HEK cell lines derived from the sub-cloning of polyclonal bulk cultures in which *CTPS1* (CTPS1-KO or CTPS1-Low) or *CTPS2* (CTPS2-KO) have been inactivated by CRISPR-Cas9 genome editing. Double CTPS1 and CTPS2-inactivated cell lines (CTPS1+CTPS2-KO) were obtained from the CTPS1-KO#5 clone. Two wild-type cell lines obtained from the sub-cloning of cells targeted by CRISPR for *CTPS1* (Ctr.#2 and #3) or control HEK cells (Ctr.#1) are also shown. (**D**) CTPS activity measured from cell lysates of control (Ctr.) or CTPS1 and/or CTPS2-deficient HEK cells (CTPS1-KO, CTPS1-Low, CTPS2-KO and CTPS1+CTPS2-KO). All cells were maintained without cytidine supplementation prior to the activity measurement, excepted for the CTPS1- and CTPS2-KO cells which were expanded in the presence of 200µM of cytidine, washed and cytidine-starved for 48h before CTPS activity was measured. Means with SEM of CTPS activity values for each group of clones are shown in the right panel. Two-tailed unpaired t-tests against Ctr. values; n.s., no significance; **, p<0.01; ****, p>0.0001. (**E**) Confluency curves as percentages (%) showing the proliferation of control (Ctr.) or CTPS1 and/or CTPS2-deficient HEK cells (CTPS1-KO, CTPS1-Low, CTPS2-KO and CTPS1+2-KO). Confluency measurement using an IncuCyte Zoom system. Cells were seeded for 24h, then treated with the indicated concentrations of cytidine. (**A-D**) Data of one representative experiment of 2 (C-D) or 3 (A, B) independent experiments. (**E**) Data of one representative experiment of 3 independent experiments.

### Inactivation of CTPS1 and/or CTPS2 in HEK cells

Thus, we decided to inactivate *CTPS1* and/or *CTPS2* by CRISPR-Cas9 genome editing in HEK cells (that express high levels of CTPS2) and Jurkat cells (that do not express CTPS1). Single RNA guides (sgRNA) targeting sequences in exons 6 and 10 of *CTPS1* or 5 and 10 of *CTPS2* were first introduced into HEK cells by transfection using a plasmid containing the sgRNA and sequences for Cas9 and puromycin resistance genes. After puromycin selection, cells were maintained in culture in the presence of cytidine to provide intracellular CTP to the cells through the salvage pathway to avoid counter selection of CTPS1- or CTPS2-deficient cells. Cytidine is a substrate of the salvage pathway and supplementation with cytidine indeed bypasses the requirement of CTPS activity for CTP synthesis [28]. Bulk cultures were first analysed by western blot to identify the most efficient guides (data not shown), then sub-cloned and clones analysed for CTPS1 and/or CTPS2 expression by western blot (**Fig. 1C**). Several clones either not expressing CTPS1 or CTPS2 (CTPS1- KO or CTPS2-KO) or expressing decreased levels (∼50%) of CTPS1 (CTPS1-low) were selected for further studies. Clones deficient for both CTPS1 and CTPS2 (CTPS1+2-null) were obtained from a CTPS1-deficient clone that was transfected with a CRISPR-Cas9 vector containing guides targeting *CTPS2*. We first analysed the CTPS enzymatic activity in cell lysates of CTPS1- and/or CTPS2-deficient cells. While the absence of CTPS2 had no significant impact on the global level of CTPS activity (**Fig. 1D**), cells expressing decreased amounts of CTPS1 showed a strongly reduced CTPS activity (∼50%) compared to controls. In cells in which CTPS1 expression was completely abrogated, CTPS activity levels were further decreased to 20% of the CTPS activity of control wild-type cells. These data indicate that the contribution of CTPS2 to the total CTPS activity of HEK cells is negligible when CTPS1 is expressed. The combined absence of CTPS1 and CTPS2 as expected completely abrogated the CTPS activity suggesting that CTPS2 likely contributes to 10-20% of the CTPS activity (although its contribution seemed to be very limited or null when CTPS1 is expressed).

We next assessed the effect of *CTPS1* and/or *CTPS2* inactivation on the proliferation of HEK cells. Growth of control, CTPS1-KO, CTPS1-low, CTPS2-KO and CTPS1+2-KO cells in the presence or absence of cytidine supplementation was evaluated using a live-cell analysis system that measures cell confluency over time. Addition of cytidine had no effect on the proliferation of control, CTPS2-deficient cells, or cells with reduced CTPS1 levels, supporting the notion that CTP production by these cell lines was sufficient to sustain cell growth (**Fig. 1E**). Although CTPS1-deficient cells were able to proliferate in the absence of cytidine supplementation, their proliferation rate/velocity was diminished as cytidine supplementation increased their cell growth (**Fig. 1E and Fig. S1**). These data indicate that the decreased CTPS activity observed in CTPS1- deficient cells impacted their ability to proliferate normally. In striking contrast, cells deficient for both CTPS1 and CTPS2 were unable to proliferate in the absence of cytidine supplementation. Cytidine supplementation restored their proliferation in a concentration-dependent manner (**Fig. 1E**). In the absence of cytidine supplementation, non-proliferating CTPS1+2-KO cells persisted in the culture for at least up to 10 days, and only after 14 days without cytidine, were a few dead cells observed accumulating in the culture (**Fig. S2*A* and B**). These cells exhibited no gross abnormalities of cell cycle (**Fig. S2C)** and intriguingly exhibited an abnormally large size (**Fig. S3**). They were able to resume proliferation as soon as cytidine was added to the medium indicating their quiescent status (**Fig. S2C**).

We then analysed the sensitivity of these different cell lines to 3-deazauridine (3-DU), a uridine analog known to be a selective inhibitor of CTPS activity, by competing with the UTP substrate [29,30]. Cells treated with different concentrations of 3-DU were evaluated for cell growth. First, we observed that the negative impact of 3-DU on proliferation was strictly due to selective inhibition of CTPS activity as supplementation of the cells with cytidine rescued cell proliferation to control levels (**Fig. 2**). Control/wild-type and CTPS2-KO cells either responded weakly or not at all to low concentrations of 3-DU, whereas proliferation of CTPS1-low and CTPS1-KO cells in response to similar concentrations of 3-DU was notably reduced or completely abolished, respectively. These results indicate that 3-DU can be used to titrate CTPS activity and suggest that CTPS2 is more prone to 3-DU inhibition than CTPS1. This could also indicate that the CTPS2 activity is lower compared to that of CTPS1 as the result of a reduced affinity for UTP (compared to CTPS1) that is more easily counteracted/displaced by the 3-DU. This also confirms that CTPS1 is the main contributor to CTPS activity in HEK cells. Taken together, these data indicate that CTPS1 is an important factor for the proliferation of HEK cells, whereas the contribution of CTPS2 is rather modest in this model, although it appears to be essential when CTPS1 is absent (e.g., in CTPS1-KO cells).

**FIGURE 2.**
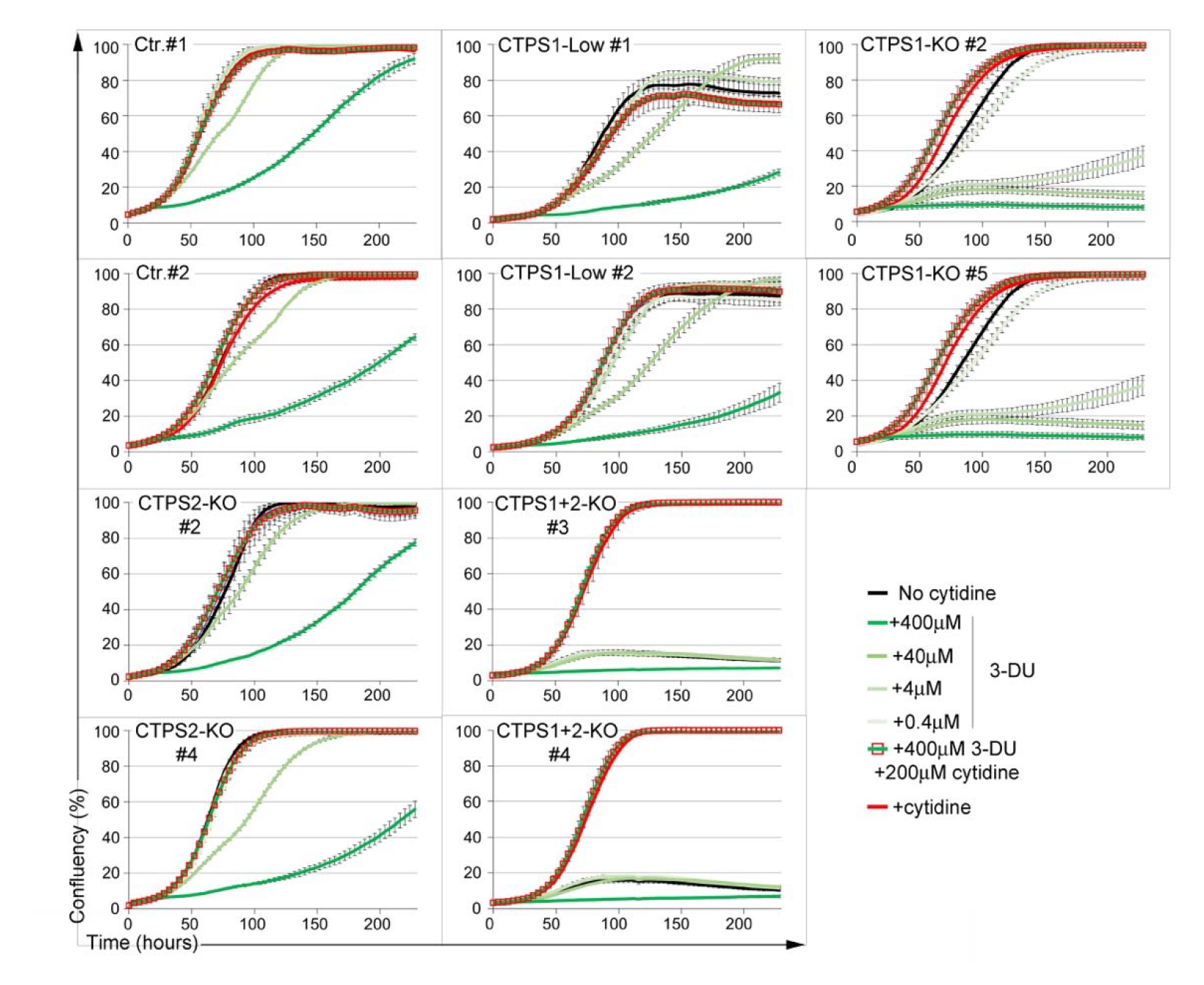
Response of CTPS1 and/or CTPS2-deficient HEK cells to 3-deazauridine. Confluency curves as percentages (%) showing the proliferation of control (Ctr.) or CTPS1 and/or CTPS2-deficient HEK cell lines (CTPS1-KO, CTPS1-Low, CTPS2-KO and CTPS1+CTPS2-KO). Confluency measurement using an IncuCyte Zoom system. Cells were seeded for 24h, then treated with the indicated concentrations of 3-deazauridine (3-DU) in the presence or not of cytidine or cytidine alone at 200µM. Data of one representative experiment of 3 independent experiments.

### Inactivation of CTPS1 in Jurkat cells

We previously reported the major role of *CTPS1* in the proliferation of activated T lymphocytes [26,27]. Multiple T lymphocyte cell lines express low levels or are negative for the expression of *CTPS2* at both RNA and protein levels, including the acute leukemia T-cell line Jurkat, in which *CTPS2* transcript and protein were undetectable (**Fig. 1**). We obtained Jurkat cells in which *CTPS1* was targeted by CRISPR-Cas9 [27]. These cells were analysed for CTPS1 expression by intracellular cytometry staining. Most of them were negative for CTPS1, in contrast to cells that were targeted with a control guide, albeit a small fraction remained positive (∼20%) (**Fig. 3A**). Cells were maintained in the presence of cytidine to avoid counter-selection of the remaining cells expressing CTPS1. Removal of cytidine supplementation after a few days led to the rapid loss of the CTPS1-deficient cells to the benefit of CTPS1-positive cells that rapidly expanded in the culture (**Fig. 3B**). The loss of CTPS1-deficient cells correlated with the appearance a massive population of cells, whose shape in FSC/SSC representation likely corresponded to either dying or dead cells (data not shown) that was further confirmed by analysing CTPS1-KO clones (see below). This suggested that the CTPS1-expressing cells had a selective growth advantage in culture, while the CTPS1-deficient cells rapidly died. Because some cells still expressed CTPS1 after inactivation by CRISPR-Cas9, we sub-cloned these cells in the presence of cytidine to support their growth. Several clones were obtained in which CTPS1 expression was completely abolished (CTPS1-KO) (**Fig. 3C**). These clones were further analyzed for cell proliferation and CTPS activity. As expected, CTPS1-KO clones had no detectable CTPS activity (**Fig. 3D**). When cytidine was removed, CTPS1-KO cells stopped proliferating (**Fig. 3E**) and were arrested in the G1 cell cycle phase as early as 24h (**Fig. 3F**). In the presence of cytidine, CTPS1-KO cells had a comparable proliferation and cell cycle progression to that of control Jurkat cells (cultured with or without cytidine). Of note, similar to what was observed with wild-type HEK cells, cytidine supplementation had no impact on the proliferation of wild-type Jurkat cells indicating that CTP was not a limiting factor for their growth (**Fig. 3E**). Further analysis of proliferation by CFSE incorporation and dilution confirmed that CTPS1-KO cells failed to proliferate in the absence of cytidine supplementation (**Fig. 3G**). Inhibiting CTPS activity in wild-type Jurkat cells by 3-DU treatment led to a strong decrease in cell proliferation that was reversed by the addition of cytidine (**Fig. 3E and G**). 3-DU treatment of Jurkat cells also resulted in a block of the G1 phase of the cell cycle (similar to that of CTPS1-KO cells in absence of cytidine) (**Fig. 3F**). The effect of CTPS1 deficiency on cell-death was further analysed in CTPS1- KO cells cultured in the presence of cytidine and then deprived of cytidine for 96 hours (**Fig. 3H** and **Fig. S4A)**. Cytidine deprivation after 48 hours led to the rapid accumulation of apoptotic (7AAD^-^ AnnexinV^+^) and dead (7AAD^+^ AnnexinV^+^) cells. Similar kinetics for the appearance of apoptotic and dead cells were observed when wild-type Jurkat cells were cultured in the presence of 3-DU. Apoptosis induced by cytidine starvation (for CTPS1-KO cells) or 3-DU treatment (for wild-type Jurkat cells) can be stopped or reduced by adding cytidine 24 h or 48 h post-deprivation, respectively (**Fig. S4B**). Taken together, these data demonstrate that CTPS1 is a key factor for Jurkat cell survival and proliferation.

**FIGURE 3.**
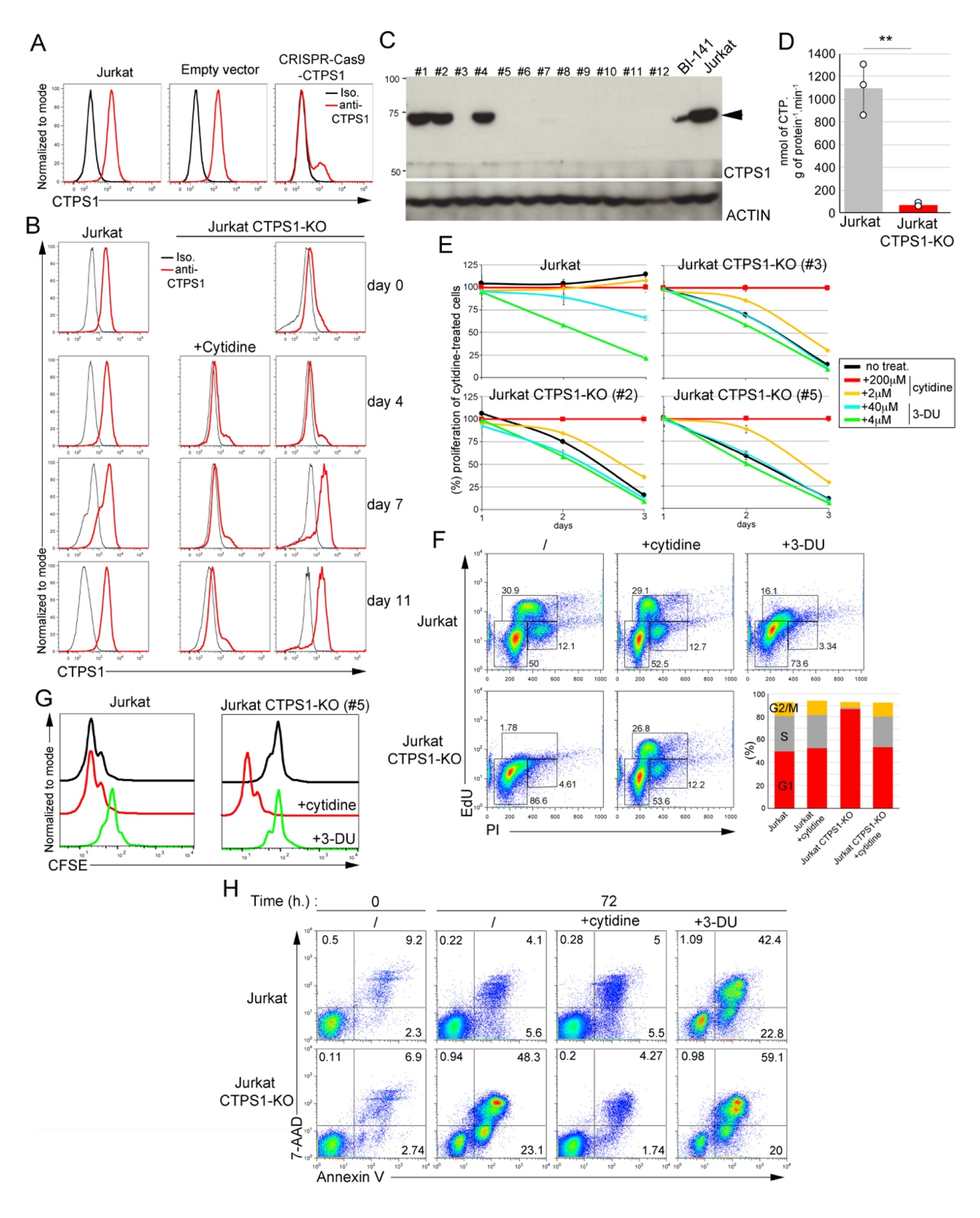
CTPS1 is required for the survival and proliferation of Jurkat cells. (**A**) Histograms from FACS analysis showing intracellular staining of CTPS1 in control Jurkat cells and Jurkat cells in which *CTPS1* has been targeted by CRISPR-Cas9 genome editing (CRISPR- Cas9-CTPS1) or with an empty vector which not contains the guide (Empty vector). Targeted cells were sorted on GFP expression and maintained in culture with cytidine before analysis. The black line corresponds to the isotype control, the red line to the anti-CTPS1 staining. (**B**) same as (**A**) except that cytidine was removed or not from the cultures for up to 11 days (**C**) Western blots of CTPS1 expression in cell lines obtained after sub-cloning of polyclonal cells shown in (**A**). ACTIN expression as a loading control. **(D)** CTPS activity measured in cell extracts of wild-type Jurkat cells and CTPS1-deficient Jurkat clones. Means with SEM of three independent experiments. Two-tailed unpaired t-test; **, p<0.01. **(E)** Proliferation graphs from Resazurin/Resofurin assays of three CTPS1-KO Jurkat cell lines (#2, #3 and #5) and wild-type Jurkat cells in the presence or not of cytidine or 3-DU at the indicated concentrations. **(F-H)** Analysis of one of the CTPS1-KO cell lines (#5) and one control cell line for cell cycle progression **(F)**, proliferation (**G**) and apoptosis (**H**) in the presence or not of 200µM cytidine or 40µM 3-DU. **(F)** FACS dot-plots of cell cycle analysis showing incorporation of EdU and IP incorporation in control or CTPS1-KO cells. Diagram on the right showing the correspondence of the gates with the G1, G2 and S phases of the cell cycle. Lower graphs showing the percentages of cells in G1, G2 and S phases from FACS data. (**G**) Histograms from FACS analysis of CFSE staining dilution-based proliferation assay. (**H**) FACS dot-plots of expression of the apoptotic/cell death annexin V and 7-AAD at 72h. Cells were seeded for 24h, and then cultured in the presence or not of 200µM cytidine or 40µM of 3-DU. (**B-H**) Jurkat correspond to Jurkat cells shown in panel A that have been transfected with an empty vector. (**A-H**) Data of one representative experiment of 3 independent experiments in (**A)**, 2 in (**B)**, 3 in (**E)** for Jurkat and Jurkat CTPS1-KO (#3), 3 in (**G)** and 3 in (**H)**.

### Differential roles of CTPS1 and CTPS2 in promoting proliferation

Our results regarding the respective role of CTPS1 and CTPS2 in the proliferation of HEK cells were in favor of a predominant role for CTPS1 in proliferation. To confirm this hypothesis, we compared the ability of CTPS2 or CTPS1 to restore the proliferation of Jurkat CTPS1-deficient cells. CTPS1-deficient cells were transduced with lentiviral vectors coding either for CTPS1 or CTPS2 along with a fluorescent mCherry reporter in the presence of cytidine and then grown in cytidine-free medium (**Fig. 4A**, left panel). Transduction with a low viral titer in which less than 1% of cells were transduced, allowed to follow the selective advantage of the transduced mCherry-positive population. From day 2, mCherry-positive cells transduced with CTPS1 rapidly expanded in culture and all cells were mCherry-positive at day 20. In contrast, expansion of mCherry-positive cells transduced with CTPS2 was delayed and only at day 15 did mCherry-positive cells begin rapidly accumulated to reach 100% at day 23. In both cultures, selection of mCherry-positive cells was associated with an initial marked decrease in the global cell viability, caused by the cytidine starvation, that recovered by day 10 in correlation with the expansion of mCherry-positive cells in the culture (**Fig. 4A**, right panel). As expected, expression of CTPS1 or CTPS2 was detectable in both cultures by western blot and was barely observed in cultures that were maintained in the presence of cytidine in which mCherry-positive cells did not expand (**Fig. 4B**). Interestingly, the mCherry staining revealed two main distinct populations in both CTPS1 or CTPS2 complemented cells (**Fig. 4C**). The subpopulation with the highest level of mCherry decreased over time for the benefit of the population with the lowest level suggesting a counter selection effect possibly due to toxicity associated with high expression of CTPS1 or CTPS2. However, the population of CTPS2-complemented cells with the lowest level which was selected over time expressed higher mCherry levels than the corresponding population in CTPS1- complemented cells (with the lowest mCherry expression).

**FIGURE 4.**
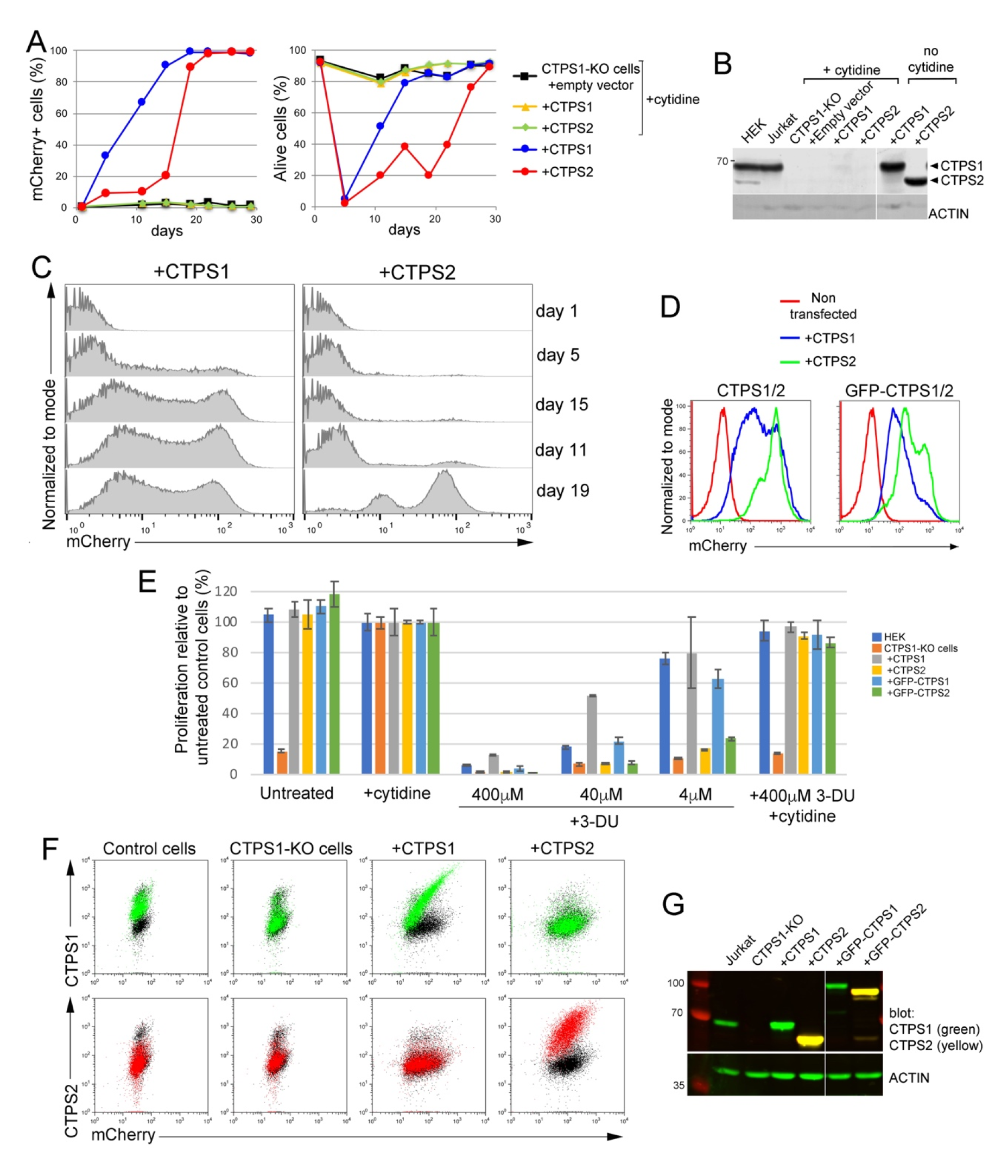
More CTPS2 than CTPS1 is required for proliferation of Jurkat cells. (**A-C)** CTPS1-deficient Jurkat cells infected with lentiviral expression vectors for CTPS1 or CTPS2 with mCherry as a reporter gene. Cells were then maintained in culture with or without cytidine. Data representative of 3 independent experiments. (**A**) Percentages (%) of mCherry-positive cells (left panel) and alive cells (right panel) based on FACS profiles. At day 0, 0,6 and 0,46 % of the cells infected with CTPS1 and CTPS2 were mCherry-positive, respectively. **(B)** Western blots for CTPS1 and CTPS2 expression in Jurkat cell lysates at day 26. (**C**) Histogram profiles from FACS analyses of mCherry expression of cells during the culture without cytidine. (**D-G**) CTPS1-deficient Jurkat cells infected with high viral titers of lentiviral vectors for CTPS1 or CTPS2 expression with mCherry as a reporter gene allowing around 94% (CTPS1/CTPS2) and 70% (GFP-CTPS1/GFP-CTPS2) of mCherry-positive cells at day 0. Cells were then maintained for 17 days in culture without cytidine for selection. (**D**) Histogram profiles from FACS analyses of mCherry expression of cells at day 55. (**E**) Bars graph of cell proliferation from resazurin/resofurin assays of the different cell lines not treated (untreated) or in the presence or not (untreated) of cytidine (200µM) or with the indicated concentrations of 3-DU. Means with SEM of experimental triplicates. (**F**) Dot-plots from FACS analysis of intracellular CTPS1 (in green) or CTPS2 (in red) and mCherry reporter expression showing that the mCherry expression is proportional to CTPS1 or CTPS2 expression. Isotype in black. (**G**) Western blots for CTPS1 and CTPS2 expression (at day 37). (**B and G**) ACTIN expression as loading control. (**A-G**) Data of one representative experiment of 3 independent experiments in **(A)**, 3 in **(C)**, 3 in **(D)** and 2 in **(E)**.

We also observed higher levels of mCherry associated with CTPS2 complementation compared to CTPS1 when CTPS1-deficient Jurkat cells were complemented using high viral titers allowing up to 90% of infected cells after a few days of selection in the absence of cytidine (in contrast to the previous experiment) (**Fig. 4D**). Similar findings were obtained when cells had been transduced with GFP tagged forms of CTPS1 and CTPS2 (denoted as GFP-CTPS1 and GFP-CTPS2). Analysis of the proliferation of the different complemented Jurkat cell populations did not reveal any differences (**Fig. 4E**). Furthermore, CTPS1-KO Jurkat cells complemented with CTPS2 forms were more sensitive to 3-DU treatment than cells complemented with CTPS1 forms, similar to our previous observations for CTPS1-deficient HEK cells that only expressed CTPS2 (CTPS1-KO cells, see **Fig. 2C**). Importantly, we showed that levels of mCherry expression directly correlated with levels of CTPS1 or CTPS2 expression analysed via intracellular staining in parallel to the mCherry expression (**Fig. 4F, G**). Therefore, taken together these data indicate that more CTPS2 than CTPS1 is required to maintain the proliferation of Jurkat cells.

Similar findings were obtained when CTPS1-deficient HEK cells (CTPS1-KO cells) and double CTPS1 and CTPS2-deficient HEK cells (CTPS1+2 KO cells) were complemented with GFP-tagged forms of CTPS1 or CTPS2 (**Fig. 5**). GFP expression was higher in cells complemented with CTPS2 compared to CTPS1 (**Fig. 5A**). Although cells with GFP-CTPS2 expressed high levels of CTPS2 (**Fig. 5A, B**), their proliferation rate/velocity appeared to be reduced compared to GFP-CTPS1-expressing cells (**Fig. 5C and Fig. S5**). As shown previously, cells complemented with CTPS2 forms were more sensitive to 3-DU treatment. Overall, the results from these two cell lines were convergent, and therefore, indicate that the role of CTPS2 is less important than that of CTPS1 to promote cell proliferation and that more CTPS2 is required to achieve the same rate of proliferation.

**FIGURE 5.**
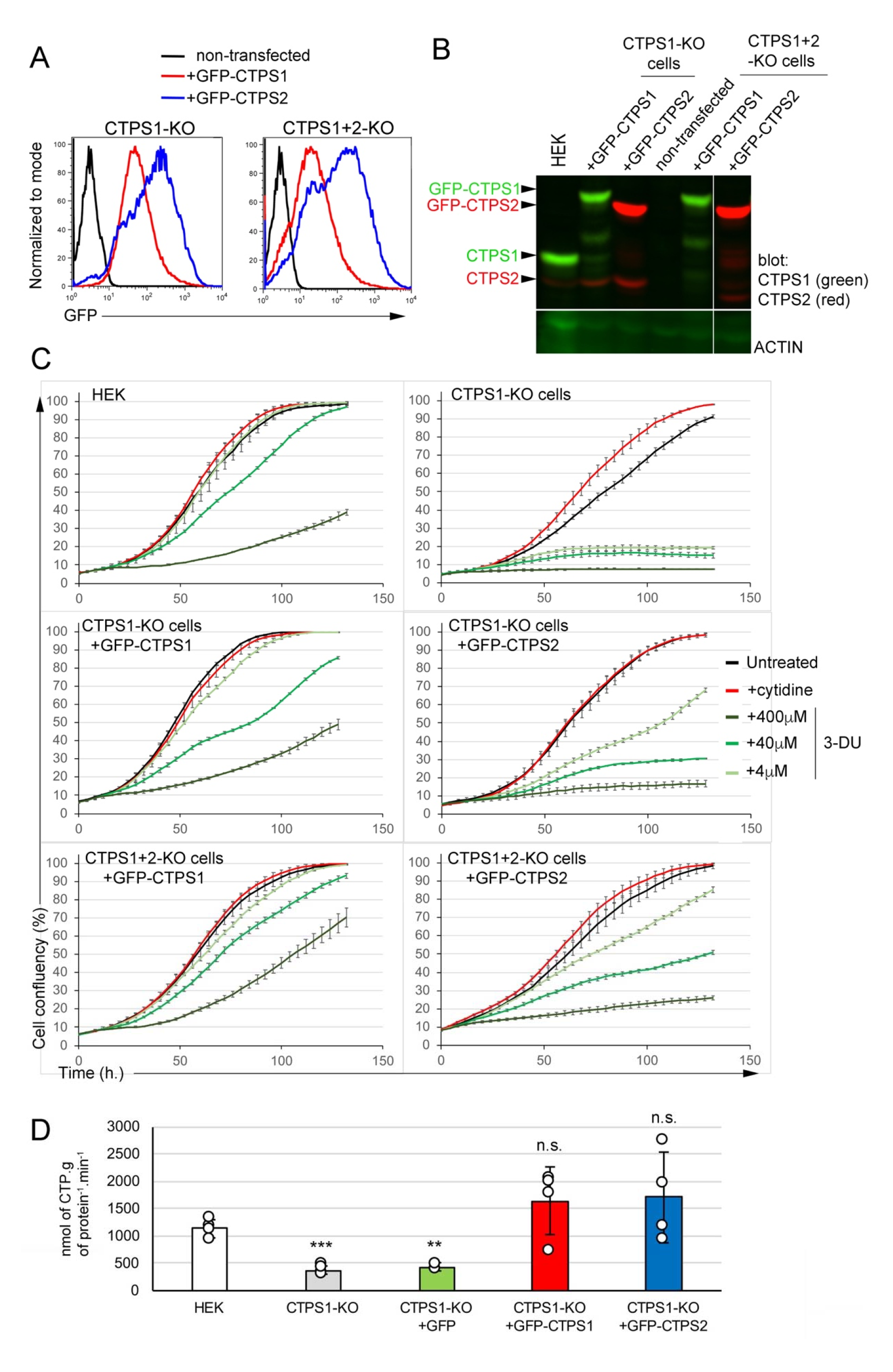
CTPS1 is more efficient than CTPS2 in restoring the proliferation of CTPS1-KO and CTPS1+2-KO HEK cells. **(A-D)** CTPS1-KO or CTPS1+2-KO cells were transfected with linearized C1 EGFP vectors containing GFP-CTPS1 or GFP-CTPS2. Cells were then maintained in culture without cytidine and sorted on GFP expression. (**A**) Histograms of GFP expression after sorting and culture in the absence of cytidine. (**B**) Western blots for CTPS1 and CTPS2 expression in cell lysates. ACTIN expression as a loading control. (**C**) Confluency curves as percentages (%) showing the proliferation. Confluency measurement using an IncuCyte Zoom system. Cells were seeded for 24h, then untreated or maintained in the presence or not of cytidine (200µM) or 3-DU with the indicated concentrations. (**D**) CTPS activity measured in cell extracts of CTPS1-KO cells reconstituted with GFP alone, GFP-CTPS1or GFP-CTPS2. Means with SEM. Data from 4 independent experiments with replicates. Two-tailed unpaired t-tests againt HEK values; n.s., no significance; **, p<0.01; ***, p>0.001. (**A-C**) Data of one representative experiment of 3 independent experiments in **(A)** and 4 in **(C)** except for CTPS1+2KO cells with GFP-CTPS2 only tested 2 times.

### Reduced enzymatic activity of recombinant CTPS2

The reduced ability of CTPS2 to sustain proliferation could be explained by lower enzymatic activity and/or differences in the regulation of its enzymatic activity as suggested by recent observations [19,31]. To test this possibility, CTPS activity was first examined in GFP- CTPS1 or GFP-CTPS2 reconstituted HEK cells and found to be roughly similar (**Fig. 5D**). This likely indicates that indeed CTPS2 enzymatic activity is weaker than that of CTPS1 as GFP- CTPS2 reconstituted cells expressed higher levels of GFP-CTPS2 (compared to GFP-CTPS1 expressing cells). In order to confirm this observation, the enzymatic activity of recombinant purified human CTPS2 and CTPS1 proteins produced in HEK cells was tested *in vitro* (**Fig. S6**). CTPS1 and CTPS2 were produced and purified using C and N-terminal tags. All CTPS2 forms exhibited reduced enzymatic activity when compared to CTPS1 (**Fig. 6A**). Interestingly, CTP was more potent at inhibiting CTPS2 than CTPS1 (**Fig. 6B**). These observations are in line with the recent observations indicating that CTPS2 contains two sites of negative feedback regulation by CTP in contrast to CTPS1 that only has one [31]. Hence, the weaker intrinsic enzymatic activity of CTPS2 likely accounts for the diminished ability of CTPS2 to support cell proliferation.

**FIGURE 6.**
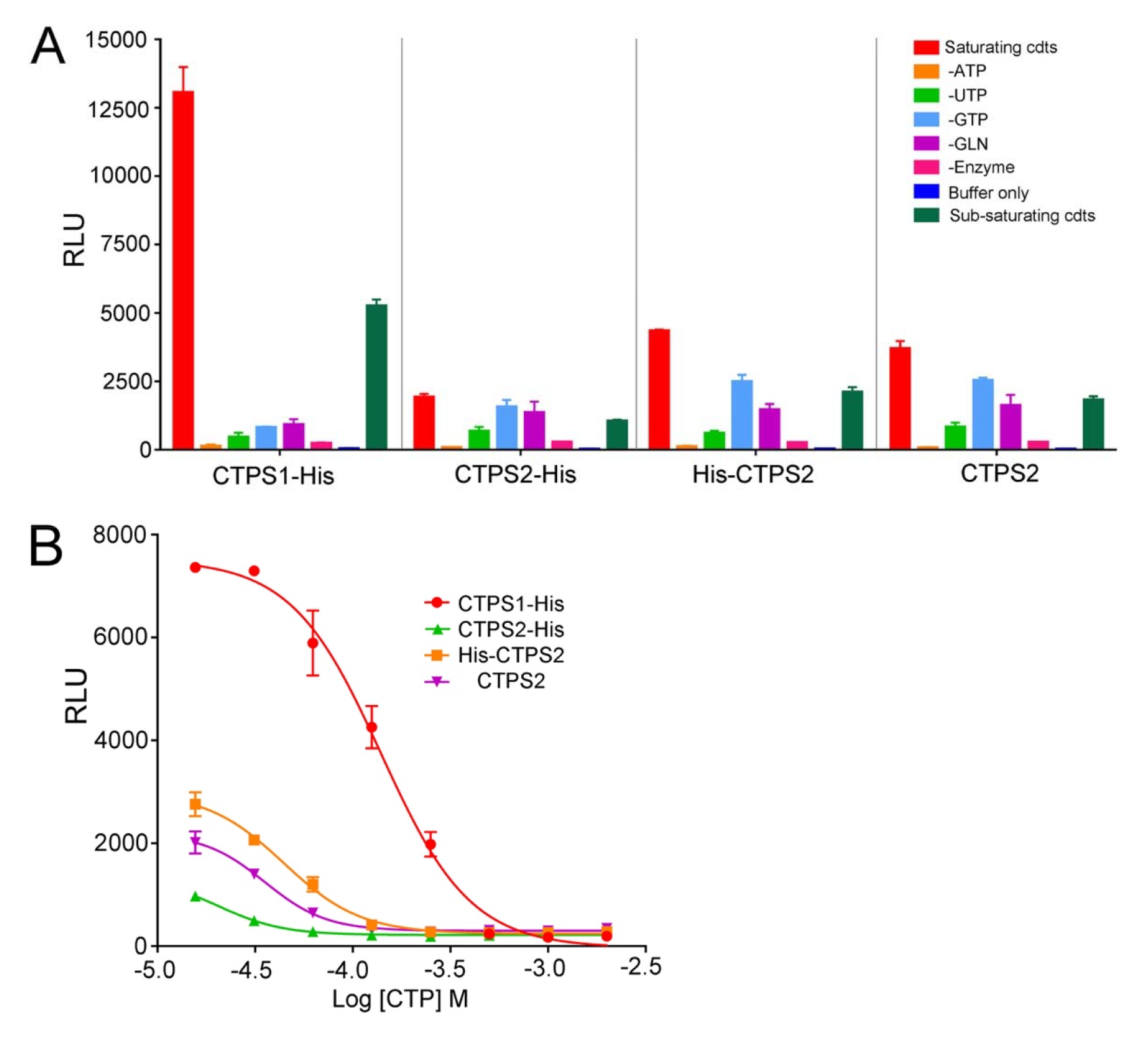
In vitro enzymatic activity of recombinant CTPS2 is lower than that CTPS1. **(A-B)** Enzymatic activity of purified recombinant tagged forms of CTPS2 and CTPS1 with C- terminal Histidine tag (CTPS1-His, CTPS2-His), CTPS2 with N-terminal Histidine tag (His-CTPS2) and CTPS2 without a tag, analyzed using ADP-Glo kinase assays (**A**) in the presence of buffer only, all substrates in saturating or sub-saturating concentrations, in conditions in which one of the substrates was removed (-ATP, -UTP,-GTP,-GLN) or without enzyme with all substrates in saturating conditions (**B**) with all substrates in saturating concentrations and with increasing concentrations of CTP. (**A-B**) Data of one representative experiment of 2 independent experiments with triplicates. Means with SEM of experimental triplicates. RLU, relative luminescence units.

### CTPS1 is essential for cancer cell growth in most cancer cell lines

To extend our observations on tumor cell lines other than HEK and Jurkat cell lines, we further examined the requirement for CTPS1 and CTPS2 for cell growth and survival of a large number of tumor cell lines from different tissues using data from the Project Achilles CRISPR-based genome-scale loss-of-function screening [32–35]. Dependency scores for *CTPS1* and *CTPS2* were extracted through the DepMap website. Interestingly, while the *CTPS2* Chronos guide depletion scores were close to 0 (mean=-0,038), highlighting its non-essentiality in cancer cell survival and proliferation, we observed a CTPS1 guide depletion (high negative scores towards - 1) in most of the 1,070 cell lines (Chronos score mean=-0,579), indicative of its essentiality (**Fig. 7**). Based on these analyses, 2 out of 1,064 cell lines (CTPS2) and 662 out of 1,070 cell lines (CTPS1) were considered to be dependent for their viability of CTPS1 or CTPS2, respectively. Taken together, these results extended our observations showing that most of the cancer cell lines tested (>1000 different cell lines) are highly dependent on CTPS1, but not or less on CTPS2 for their survival and proliferation.

**FIGURE 7.**
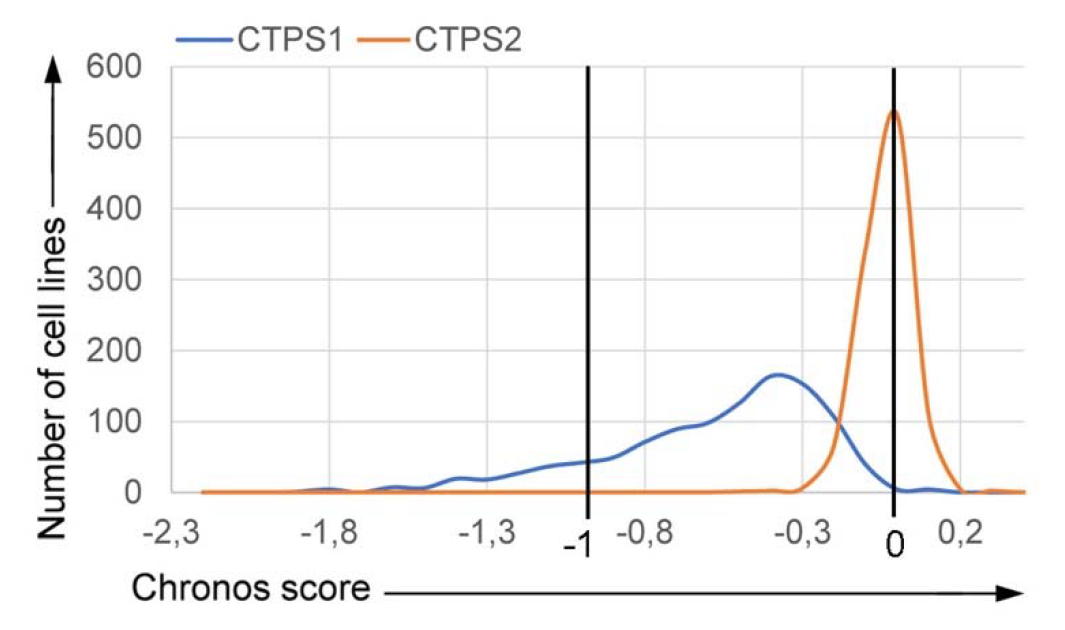
CTPS1 is an essential gene for survival and proliferation of most cancer cell lines. (**A**) Chronos score of 1064 cell lines for CTPS1 and CTPS2, extracted from the DepMap 22Q1 Public+Score Chronos dataset. Counterselection of CRISPR-targeted cells leads to guide depletion and is translated by a negative score. A score of 0 corresponds to a non-essential gene, while a negative score shows a dependency for the concerned gene. The score of −1 corresponds to the median of all essential genes from the DepMap database. For this DepMap release: DepMap, Broad (2022): DepMap 22Q1 Public. https://doi.org/10.6084/m9.figshare.19139906.v1.

## DISCUSSION

Our results in HEK and Jurkat cell lines models indicate that CTPS1 and CTPS2 are partially redundant for cell proliferation. The role of CTPS2 in proliferation is less important than that of CTPS1. Our results indicate that higher levels of CTPS2 than CTPS1 are necessary to achieve similar proliferation rates. This could be explained by the lower intrinsic enzymatic activity of CTPS2, although we cannot formally exclude other mechanisms independent of the intrinsic enzymatic activity. Along these lines, we previously showed that CTPS1 was critical to sustain proliferation of activated T lymphocytes, despite their expression of CTPS2. Thus, in activated T lymphocytes CTPS2 does not compensate for the defect in CTPS1, indicating that the role of CTPS2 to promote proliferation is minimal. Further in support of differential roles of CTPS1 and CTPS2, recent studies showed that CTPS2 enzymatic regulation is different than that of CTPS1. In contrast to CTPS1, CTPS2 contains two sites of negative feedback regulation by CTP that might contribute to its lower activity by being more sensitive to CTP and thus more rapidly inhibited than CTPS1[31]. Our results also indicate that CTPS2 is more prone to inhibition by 3- DU, a competitive inhibitor of UTP that very likely reflects a low affinity to UTP, thus leading to a lower intrinsic enzymatic activity (compared to CTPS1). Consequently, both specific regulatory and intrinsic determinants account for the low activity of CTPS2. Altogether, the higher activity of CTPS1 may be required *in vivo* to rapidly meet a critical need of CTP in highly proliferating cells, like activated T lymphocytes or cancer cells, while CTPS2, which has a lower activity, may be rather involved in maintaining basal cellular CTP pools in general.

Interestingly, full depletion of the CTPS activity (by inactivation of CTPS1 and CTPS2) has distinct consequences in Jurkat versus HEK cells, although proliferation is completely abrogated in both cell lines following CTPS inactivation. Jurkat cells without CTPS1 are blocked in the G1 phase of the cell cycle (if no CTP is provided to compensate for the lack of CTPS activity) and rapidly die by apoptosis within the next 48 hours. In contrast, HEK cells deficient for both CTPS1 and CTPS2 stopped growing in the absence of cytidine supplementation, but only a small fraction of cells died even after 10 days of CTP deprivation. When CTP was added back into the medium, cells rapidly recovered and proliferated even after 14 days of deprivation. These results indicate that HEK cells in the absence of CTPS activity or CTP are resistant to apoptosis and become quiescent (defined as a reversible non-proliferating state). Such behavior might be explained by the embryonic origin of HEK cells and by their capacity to survive under various selective conditions [36]. Deprivation of cellular CTP likely results in stalled replication forks, leading to replicative stress, a complex phenomenon that occurs when replication is impeded [37]. Depending on the nature of the replicative defect or the source of the replicative stress, the response can involve different mechanisms and consequences including apoptosis. However, the effect of CTP deprivation or defective CTPS activity on replication stress response is currently unknown and warrants further study. We also noticed that the size of HEK cells deficient for both CTPS1 and CTPS2 markedly increased upon CTP deprivation. Jurkat cells treated with 3-DU, CTPS1-deficient Jurkat cells and CTPS1-deficient T cells of patients also exhibited a larger size compared to their normal size (Minet N., Martin E. and S. Latour, unpublished observations). We have no clear explanation for this phenomenon.

Our results are important in the context of the development of CTPS1 inhibitors. We previously proposed that CTPS1 could represent a therapeutic target to suppress adaptive immune responses by specifically blocking the expansion of T lymphocytes and specific inhibitors of CTPS1 have been recently obtained that block the proliferation of Jurkat cells and human primary T cells [31]. Since high levels of CTPS activity have been reported in cancer cells [20,21], in particular in lymphomas [22], inhibition of CTPS1 represents a potent therapeutic approach to block cancer progression. Our analysis of a large number of tumour cell lines (>1000) from the Project Achilles CRISPR-based genome-scale loss-of-function screening and gene copy numbers from CCLE data confirms that *CTPS1* is an essential factor in cell growth for most of these cell lines. Furthermore, our experimental results with HEK and Jurkat cell lines suggest that in tumor cells expressing CTPS1 only or in association with a low expression of CTPS2 (like Jurkat cells), inhibition of CTPS1 could lead to the rapid death of the cells by apoptosis. However, in cells co-expressing both CTPS1 and CTPS2, inhibition of CTPS1 would be not sufficient to block proliferation as CTPS1-deficient HEK cells can still proliferate (although the rate of proliferation was diminished). The contribution of CTPS2 to proliferation thus appears to be essential when CTPS1 is absent, while minimal when CTPS1 is present. Thus, the efficacy of a treatment targeting CTPS1 might be particularly dependent on the level of CTPS2 in cancer cells.

Interestingly, among the different tumor cell lines of hematopoietic origin we tested, T-cell lymphomas appear to be those in which the expression of CTPS2 is the lowest or undetectable. The myeloid cell line U937 also shows a weak expression of CTPS2. In contrast, B-cell lymphomas expressed more substantial levels of CTPS2, and a recent study suggested that the inactivation of both CTPS1 and CTPS2 is required to fully block Epstein Barr virus-driven B cell proliferation [38]. Thus, neoplasms of T-cell origin, for which there is an important unmet medical need may represent a primary indication for CTPS1 inhibition.

Finally, as we previously observed for CTPS1-deficient primary T cells (14, 15), supplementation of CTPS-deficient HEK and Jurkat cells with cytidine through the salvage pathway restores proliferation. Hence, delivery of cytidine (or of a CTP precursor) could mitigate unwanted CTP deprivation-induced cell death or immunosuppression (in the case of cancer treatment) that might result from the use of therapeutic CTPS inhibitors.

In conclusion, our study is an important contribution to the understanding of the respective roles of CTPS1 and CTPS2 in cell proliferation in particular in cancer cells, for which limited information is currently available.

## MATERIALS AND METHODS

### Plasmids

Single-guide RNAs (sgRNAs) targeting exons 6 or 10 of *CTPS1* or exons 5 and 10 of *CTPS2* were designed as previously described [39] and cloned in the LentiCRISPR V1 (pXPR_001) or pSpCas9(BB)-2A-GFP (PX458) (Addgene plasmid 48138) vectors. The sgRNA sequences were as follows: CTPS1 exon 6 F: CACCGAGTGTTCGGGAACTTAG / R: AAACCTAAGTTCCCGAACACTC and 10 F: CACCGGCTTCGTGGTAGCGCAC / R: AAACGTGCGCTACCACGAAGCC; CTPS2 exon 5 F: CACCGCGAAGGAATGCCGTTTG / R: AAACCAAACGGCATTCCTTCG and 10 F: CACCGGAAGATCACTGAAACCG / R AAACCGGTTTCAGTGATCTTC. Extinction efficiency was verified by western blot or intracellular staining of puromycin-selected or GFP-sorted cell bulks before subcloning. For complementation experiments, full-length CTPS1 and CTPS2 cDNAs were obtained by PCR as previously described [26] using the Q5® High-Fidelity DNA Polymerase (New England Biolabs). For CTPS1, forward 5′-AAGCAGACTAGTCCACCATGAAGTACATTCTGG-3′ and reverse 5′- AAGCAGGCGGCCGCTCAGTCATGATTTATTGATGGAAACTTC-3′ were used. For CTPS2, forward 5′-AAGCAGACTAGTCCACCATGAAGTACATCCTG-3′ and reverse 5′- AAGCAGGCGGCCGCTCAGCTTATTTCCAACTCAGC-3′ were used. The cDNAs were verified by sequencing and inserted into a bicistronic lentiviral expression vector encoding mCherry as a reporter (pLVX-EF1α-IRES-mCherry Vector, Clontech cat#631987). For GFP tagging, the cDNAs were inserted into a pEGFP-C1 vector (Clontech cat#6084-1) using the following primers. For CTPS1, forward 5′-AAGCAGGGTACCCCACCATGAAGTACATTCTGG-3′ and reverse 5′- AAGCAGGGATCCTCAGTCATGATTTATTGATGGAAACTTC-3′; for CTPS2, forward 5′- AAGCAGGGTACCCCACCATGAAGTACATCCTG-3′ and reverse 5′-AAGCAGGGATCCTCAGCTTATTTCCAACTCAGC-3′ were used. All constructs were validated by Sanger sequencing using the BigDye™ Terminator v3.1 Cycle Sequencing Kit (Life Technologies) and a 3500xL Genetic Analyzer (Applied Biosystems) according to the manufacturer’s instructions. Sequence analysis was performed using DNADynamo (BlueTractorSoftware).

### CTPS1 and CTPS2 gene expression analysis

Total RNA was isolated from cell lines using the RNeasy Mini kit (Qiagen) and reverse transcription was performed using Superscript II First Strand Synthesis System (Invitrogen). cDNAs were used as a template for CTPS1 and CTPS2 gene expression by RT–qPCR.Gene expression assays were performedwith Assayson-Demand probe and primer combinations (CTPS1, Hs01041858; CTPS2, Hs00219845; GAPDH, Hs027558991) from Applied Biosystem labelled with 6-carboxy-fluorescein (FAM) dye, and universal reaction mixture. Real time quantitative PCRs for GAPDH, CTPS1 and CTPS2 were performed in triplicate using a LightCycler VIIA7 System (Roche). Expression levels were determined by relative quantification using the comparative threshold cycle method 2DDCt in which DDCt is determined as follows: (Ct^target^ ^gene^- Ct^reference^ ^gene^) target tissue - (Ct^target^ ^gene^ - Ct^reference^ ^gene^) calibrator tissue. The results shown in arbitrary units (a.u.) have been normalized to GAPDH gene expression (reference gene) and are presented as the relative change in gene expression normalized against the calibrator sample corresponding to HEK cells.

### Cell culture

Jurkat cells (RRID:CVCL_0065, ATCC) were cultured in RPMI, 10% fetal bovine serum (FBS), 1% penicillin-streptomycin (PS) (complete RPMI). HEK 293T cells (RRID:CVCL_0063, ATCC) were cultured in DMEM, 10% FBS, 1% PS (complete DMEM). Calcium and magnesium-free PBS and 1x trypsin-EDTA were respectively used to wash the cells and to detach adherent cells. All of the abovementioned reagents were from Life Technologies. CTPS1 or/and CTPS2-deficient cell lines were maintained with 200µM of cytidine (Sigma Aldrich). 3-deazauridine (3-DU) (Sigma Aldrich) was used as a non-selective inhibitor of CTPS1 and CTPS2.

### Cell transfection and transduction

HEK 293T cells were transfected by electroporation using the Gene Pulser Xcell (Bio-Rad) or by lipofection with Lipofectamine 2000 (Life Technologies). pEGFP-C1 plasmids with CTPS1 or CTPS2 were linearized before electroporation, purified on High Pure PCR Product Purification Kit columns (Roche) and eluted in water. After electroporation, cells having incorporated the vector were selected using puromycin (Invivogen) for the pXPR_001 vector, sorted based on GFP expression for the PX458 vector and the pEGFP-C1 vector, or on mCherry expression for the pLVX vector (RRID:Addgene_174088) using cell sorter (Sony SH800). For low efficiency transductions, Jurkat CTPS1-KO cells were infected lentiviral particles containing lentiviral vectors for CTPS1 or CTPS2 expression as previously described (Martin et al., 2020). For high efficiency transductions, viral particles produced at the VVTG platform at Hospital Necker (VVTG platform, SFR Necker, Paris, France) were concentrated by ultracentrifugation and the cells were infected with a MOI of 10.

For CRISPR-Cas9-mediated gene inactivation, Jurkat cells were transfected by electroporation using the Nepa21 electroporator (Nepagene) with PX458 plasmids, sorted on EGFP expression, and subcloned as previously described (Martin et al., 2020). HEK 293T cells were electroporated with LentiCRISPR V1 plasmids, selected using puromycin and subcloned. HEK cells in which both CTPS1 and CTPS2 were inactivated were obtained by inactivation of CTPS2 using the same approach on one the CTPS1-deficient HEK cell line obtained (CTPS1-KO clone#5). Cells were maintained with 200µM of cytidine after electroporation to avoid counter-selection.

### Cell proliferation

For the resazurin proliferation assay, cells were seeded at a density of 150000 cells/ml in 96-well plates. At the 24 and 48 hours, 10µL of cell-titer-blue (Promega) were added and cells further incubated at 37°C for up to 4 hours. Absorbance at 560/595nm was measured and analysed according to the manufacturer’s instructions using a Tecan Infinite 200 Pro-plate reader (Tecan Life Sciences). For CSFE incorporation based proliferation assay, cells were incubated overnight with 3mM of hydroxyurea (Sigma), washed and labelled with CFSE (Invitrogen) according to the manufacturer’s instructions, resuspended in complete medium and cultured for 4 days. Cells were then analysed using a LSRFortessa X-20 cytometer and the FlowJo software (BD Biosciences). Proliferation of HEK cells was analysed as a percentage of confluency. For that, cells were seeded in complete culture medium at a density of 6250 cells/cm² for 24h and placed in an IncuCyte Zoom Live-Cell analysis system (Sartorius). Phase contrast and GFP fluorescence pictures were taken every 3 hours and analysed using a confluency mask.

### Cell cycle

Cells were washed, resuspended in complete medium and cultured for 24 hours with the indicated concentrations of cytidine or 3-DU. The cells were then incubated for a further 1 hour with 10µM EdU, fixed and stained according to the manufacturer’s instructions (Click-iT™ EdU Cell Proliferation Kit for Imaging, Life Technologies). All data were collected on LSR Fortessa X-20 cytometer and data were analysed using FlowJo software (RRID:SCR_008520, BD Biosciences)

### Annexin V/7-AAD apoptosis

Cells were washed and seeded at a density of 150 000 cells/mL in the presence or absence of 40µM of 3-deazauridine or 200µM of cytidine in 24 well plates (one well per time point). At each time point, the cells were stained with FITC-conjugated Annexin V and 7-AAD according to the manufacturer’s instructions (BD Biosciences) and analysed using a LSRFortessa X-20 cytometer and FlowJo software (RRID:SCR_008520, BD Biosciences).

### Intracellular FACS

The cells were fixed and permeabilized using the IntraPrep Permeabilization Reagent (Beckman Coulter A07803) according to the manufacturer’s instructions. Cells were stained first with an anti-CTPS1 antibody (Abcam ab133743), an anti-CTPS2 antibody (Abcam ab190462) or an isotype-matched antibody (rabbit IgG, Abcam ab172730) and then labeled with an AF647-goat anti-rabbit secondary antibody. All data were collected on an LSRFortessa cytometer and analysed using Flow-Jo version 9.3.2 software (RRID:SCR_008520, BD Biosciences).

### Western Blot

Cell lysates were prepared and analysed using standard procedures as previously described [26,27]. The following antibodies were used for immunoblotting: rabbit polyclonal anti-actin (Sigma cat# A2066), rabbit monoclonal anti-CTPS1 (Abcam ab133743), rabbit polyclonal anti-CTPS2 (Abcam ab190462), rabbit anti-GFP (Cell Signaling 2956S). The following antibodies were used for detection: IRDye 680RD or IRDye 800CW-conjugated goat anti-rabbit (Li-cor 925- 68071, Li-cor 925-32211) or goat anti-mouse (Li-cor 925-68070 and Li-cor 925-32210) and membranes analyzed with the Odyssey CLX imager (Li-Cor).

### CTPS activity measurement

CTPS activity was measured in cell extracts as previously described [26,27,40]. Briefly, cell pellets were sonicated on ice. Thirty micrograms of proteins were used for measuring CTPS activity in reaction mixture containing 1.7 mM EDTA, 13 mM MgCl2, 1.3 mM ATP, 0.2 mM GTP, 13 mM glutamine, 1.3 mM phosphoenolpyruvate, 10 mM NaF, 1.3 mM UTP, and 10 μM stable CTP isotope as an internal standard in 10 μL HEPES buffer, 70 mM, at pH 8.0. The reaction mixtures were incubated at 37 C° for 90 minutes, and then the enzymatic reaction was stopped by the addition of 2 volumes of HClO4. Five microliters of sample were injected onto an Acquity HSS T3 column, 1.8-μm particle size, 2.1 Å∼ 100 mm (Waters), connected to an Acquity H-Class ultra-performance liquid chromatography interfaced with a Xevo TQ-S triple-quadrupole mass spectrometer (Waters), both controlled by MassLynx software (MassLynx, RRID:SCR_014271, Waters). The CTP identification and detection were performed in the electrospray positive ion mode with multiple reaction monitoring mode (MRM). Quantification was performed using TargetLynx software. The threshold of CTPS activity detection corresponds to the mean of activity detected in Jurkat cells deficient for CTPS1 (that did not express CTPS2) from 4 independent experiments (51±5 nmol of CTP/g protein/min) (27).

### CTPS activity on purified enzymes

The purified N-terminal His_6_-tag with extended linker form containing a cleavage site for the TEV protease (underlined) CTPS2 form (MDHHHHHHDTT<u>ENLYFQ</u>GGSGS-CTPS1), and the C-terminal FLAG-His_8_-tag CTPS1 and CTPS2 forms (CTPS1/CTPS2- GGDYKDDDDKGGHHHHHHHH) were provided by Proteros (Martinsried, Germany). Briefly, tagged proteins were produced in HEK cells and purified by Ni affinity chromatography. CTPS2 without tag was obtained by digestion of the His-CTPS2 with the TEV protease and purified by size exclusion chromatography. Purity of the proteins was >90% after verification by peptide finger mass print analysis and by analyzing 5μg of proteins by SDS-PAGE on 10% acrylamide gels revealed by Coomassie blue staining (as shown **in Supplemental Figure 6**). CTPS activity of purified CTPS1 and CTPS2 proteins was then analyzed using ADP Glo Kinase assay (Promega). CTPS1 and CTPS2 were diluted in assay buffer (50 mM Trizma, 10 mM MgCl2, 2mM L-cysteine, 0.01% Tween-20, pH 8.0) and enzyme reaction initiated by addition of substrates and co-factors at sub-saturating concentrations (0.31 mM UltraPure ATP, 0.034 mM GTP, 0.48 mM UTP, 0.19 mM L-glutamine for CTPS1 or 0.35 mM UltraPure ATP, 0.025 mM GTP, 0.5 mM UTP, 0.06 mM L-glutamine for CTPS2; final enzyme concentration 0.8 µM). Sub-saturating concentrations were determined per isoform by performing individual substrate titrations against a mastermix of remaining key substrates and co-factors held at the same saturating concentrations for each isoform (2mM UltraPure ATP, 0.1mM GTP, 2mM UTP, 2mM L-glutamine) [41]. Of note, the different activity assay conditions which vary slightly in reactant concentrations for CTPS1 and CTPS2 were determined across multiple experiments. After incubation for 50 minutes at 20°C (within the linear phase of reaction) the reaction was terminated, and ADP product formation quantified using the ADP-Glo^TM^ Max assay system (Promega) according to the manufacturer’s instructions. Michaelis-Menten plots were subsequently determined per substrate and co-factor over multiple repeat experiments to derive final activity assay concentrations.

### Analysis of the requirement for CTPS1 and CTPS2 in cancer cells

The dependency scores for CTPS1 and CTPS2 from the Project Achilles CRISPR-based genome-scale loss-of-function screening were downloaded through the Cancer Dependency Map website (https://depmap.org/) using the latest available data (DepMap 22Q1 Public+Score Chronos dataset).

## DATA AND REAGENT AVAILABILITY STATEMENT

Datasets and reagents generated during the current study are available from the corresponding author on reasonable request.

DepMap 22Q1 public data are available on https://doi.org/10.6084/m9.figshare.19139906.v1.

## ACKNOWLEDGMENTS

We acknowledge Meriem Rahmani, the cytometry platform (SFR Necker, Paris France) directed by Corinne Cordier, the cellular imaging platform (Imagine Institute, Paris, France) directed by Meriem Garfa-Traoré for their technical assistance. We thank Andrew Parker (present Chief Executive Officer of Step-Pharma) for discussions and the critical reading of the manuscript. We also thank the members of the Latour lab for discussions. This work was supported by grants from Ligue Contre le Cancer - Equipe Labellisée (France) (to SL), Inserm (France), ANR-18- CE15-0025-01 (to SL) and ANR-10-IAHU- 01 (Imagine Institute) and Proof of Concept ERC- 2015-PoC_Master/ERC-2015-PoC_680465_SAFEIMMUNOSUPPRESS (to AF, SL). NM is a fellowship recipient of the Association Nationale de la Recherche Technologique (ANRT) under agreement with the industrial partner Step Pharma. SL is a senior scientist at the Centre National de la Recherche Scientifique (CNRS, France).

## AUTHOR CONTRIBUTIONS

NM designed and performed experiments and analyzed the data. NM and SL wrote the manuscript. A-CB, RM performed experiments and analyzed data. HA, CS, TB, EM, SL, RM, DL, PB and AF analyzed, interpreted and reviewed data. SL designed and supervised the research.

## Supplementary Material for

Differential roles of CTP synthetases CTPS1 and CTPS2 in cell proliferation inet et al.

---

**FIGURE S1.**
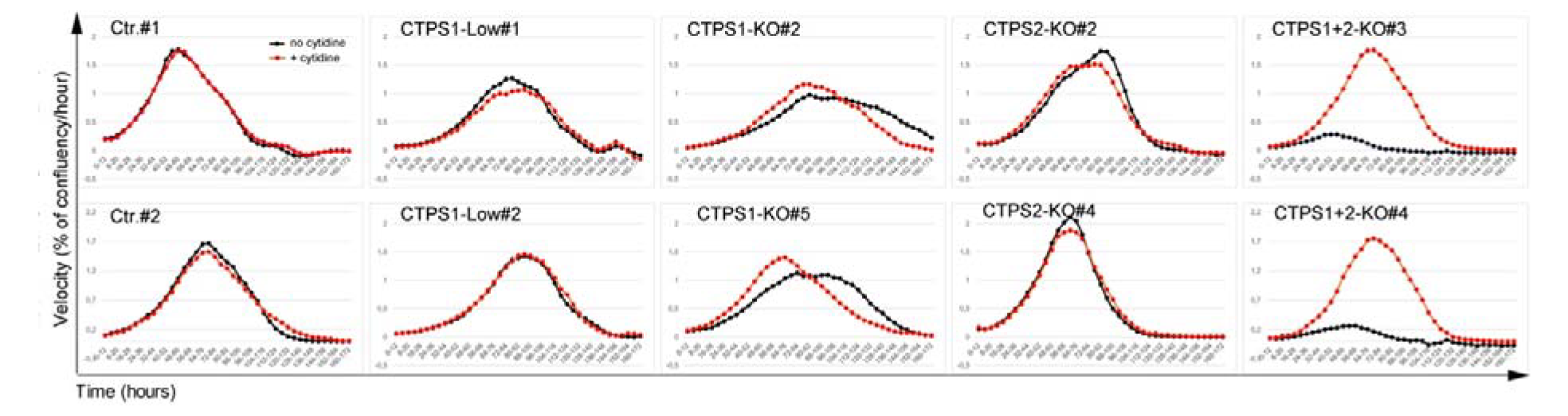
Different contributions of CTPS1 and CTPS2 to proliferation of HEK cells. Proliferation velocity, corresponding to the Figure 1E. Velocity for each population in presence (red dotted ne) or absence (black dotted line) of cytidine supplementation (200µM), corresponding to the increase in onfluency per hour on 12-hour periods. Data of one representative experiment of 3 independent xperiments.

**FIGURE S2.**
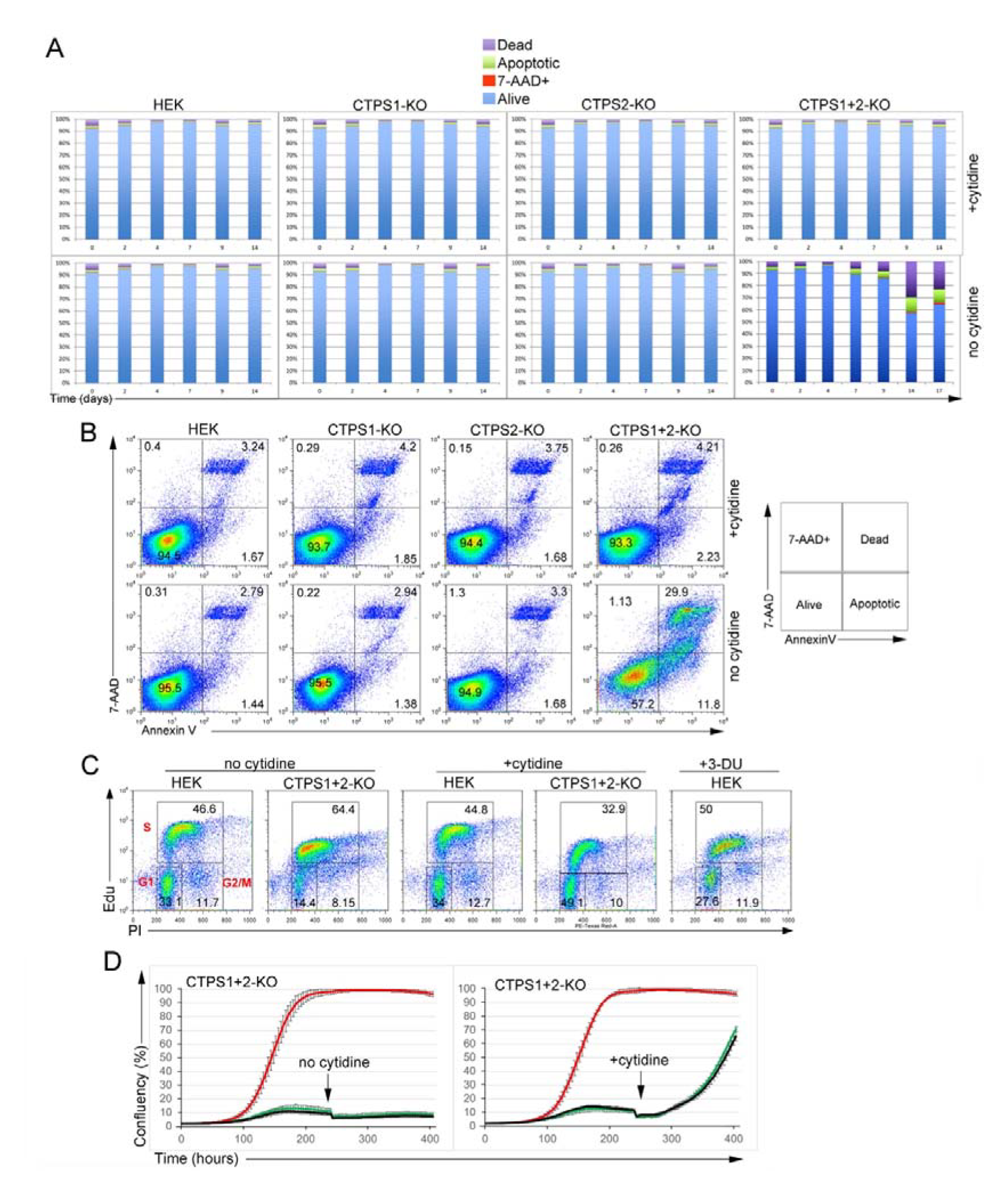
Inactivation of CTPS1 and CTPS2 in HEK cells induces quiescent state cell cycle with a oderate level of cell death. (**A-B**) Analysis of apoptotic/cell death markers of CTPS1 and/or CTPS2-deficient HEK cell lines (CTPS1-KO, TPS2-KO and CTPS1+CTPS2-KO). Cells were seeded for 24h, and then cultured in the presence or not of 00μM cytidine. Apoptotic/cell death markers annexin V and 7-AAD analyzed by flow cytometry at day 2, 4, 7, 9, 14 and 17 of culture (**A**) FACS dot-plots of annexin V and 7-AAD stainings at day 14. On the right, orrespondence between staining gates and cell phenotypes (B) Bar graphs from flow-cytometry data as resented in (A) showing the percentage (%) of dead, 7-AAD+, alive and apoptotic cells. **C)** FACS dot-plots of cell cycle analysis showing staining of EdU and PI incorporation in control HEK or TPS1+2-KO cells treated or not with 200µM cytidine or 40µM 3-DU as indicated. Gates corresponding to 1, G2/M and S phases of the cell cycle are indicated in red in the first left dot-plot. Lower graphs showing e percentages of cells in G1, G2 and S phases from FACS data. **D)** Confluency curves as percentages (%) showing the proliferation of HEK cells deficient for both CTPS1 nd CTPS2 (CTPS1+CTPS2-KO). Confluency measurement using an IncuCyte Zoom system. Cells were eeded for 24h, then cultured in the presence or not (black line) of 200µM of cytidine (red line) or 40µM of 3-U (green line). The black arrows indicate the time at which cytidine was added. (**A-C**) Data of one presentative experiment of 3 independent experiments.

**FIGURE S3.**
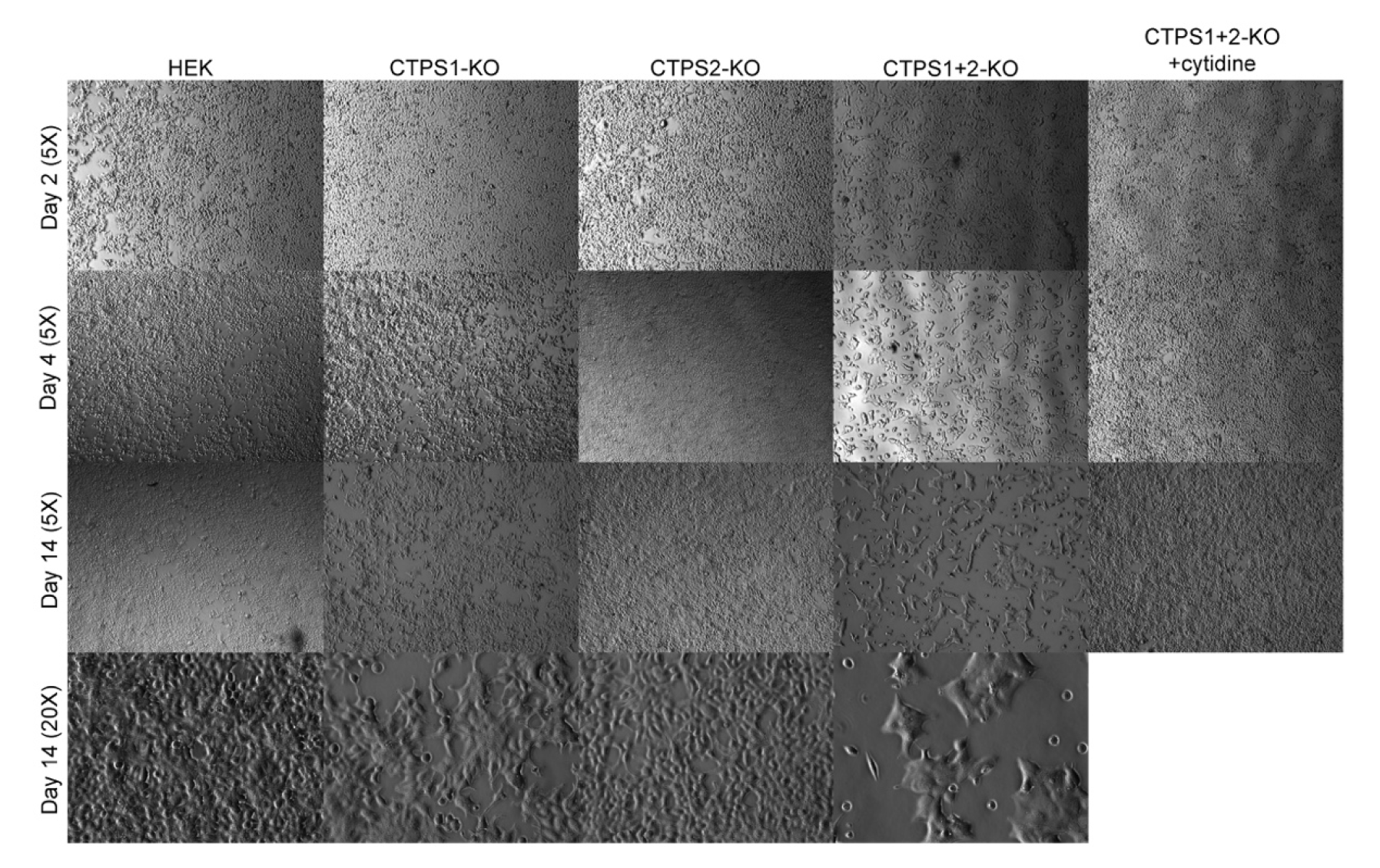
Inactivation of CTPS1 and CTPS2 in HEK cells induces an increase in cell size. (**C**) Analysis of cell confluency of CTPS1 and/or CTPS2-deficient HEK cell lines (CTPS1-KO, CTPS2-KO and TPS1+CTPS2-KO). Cells were seeded for 24h, and then cultured in the presence or not of 200μM cytidine. ictures taken with a Zeiss Axio Vert A1 at day 2, 4, 7, 9, 14 and 17 of culture. Data of one representative xperiment of 2 independent experiments except that day 17 was only tested one time.

**FIGURE S4.**
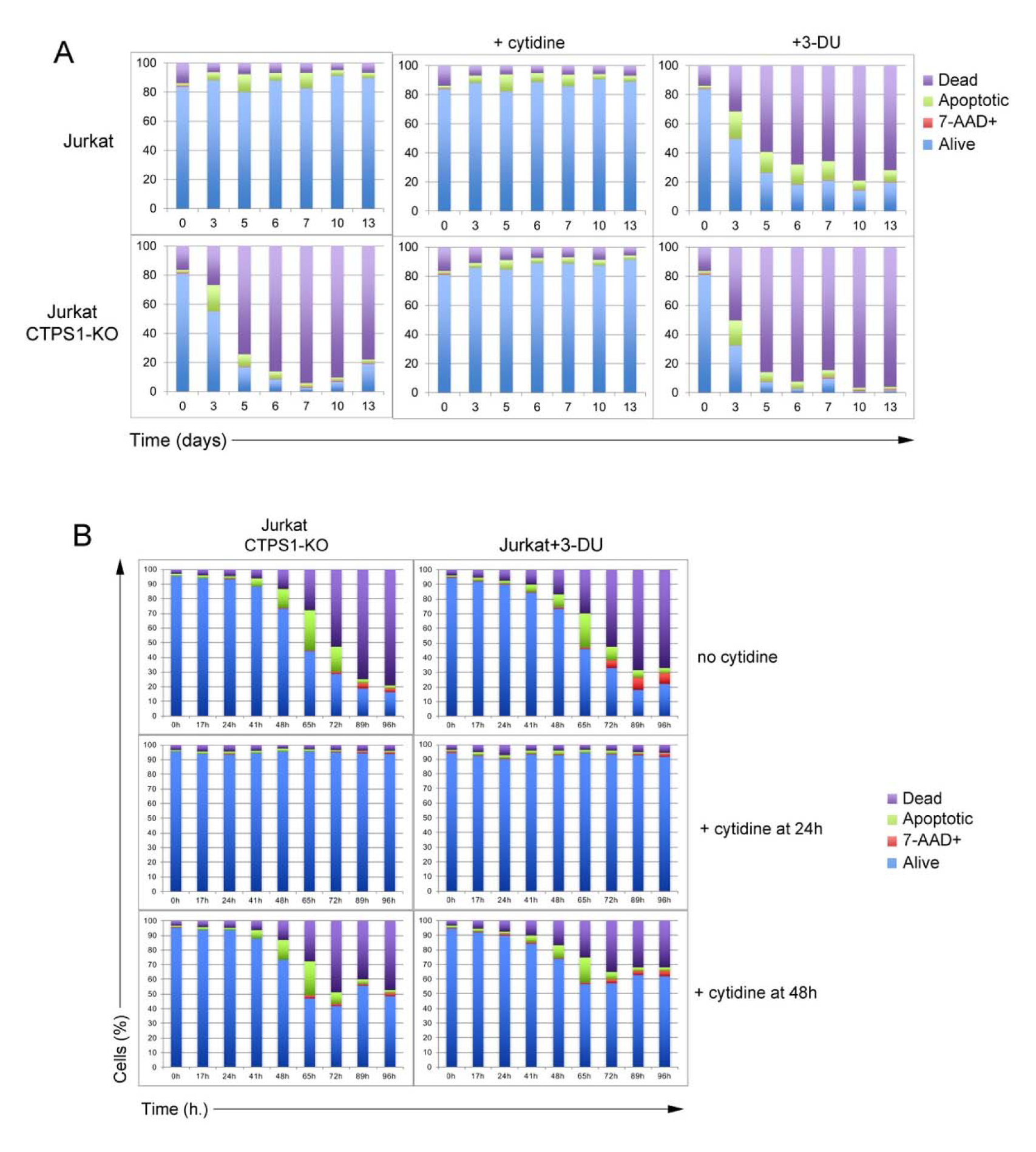
Inactivation of CTPS1 in Jurkat cells results in rapid cell death and can be counteracted by supplementation of cytidine at 24 hours but not at 48 hours. **(A, B)** Cells were seeded for 24h, and then cultured in the presence or not of 200µM of cytidine or 40µM of 3-DU. Apoptotic/cell death markers annexin V and 7-AAD were analyzed by flow cytometry at 3, 5, 6, 7, 65, 10 and 13 days. Bar graphs from flow-cytometry data as presented in Figure 4 panel (H) showing the percentage (%) of dead, 7-AAD+, alive and apoptotic cells at 17, 24, 41, 48, 65, 72, 89 and 96 hours (h.). **(B)** Same as (A) except that 200µM of cytidine has been added at 24h or 48h. **(A, B)** Data of one representative experiment of 3 (A) or 2 (B) independent experiments.

**FIGURE S5.**
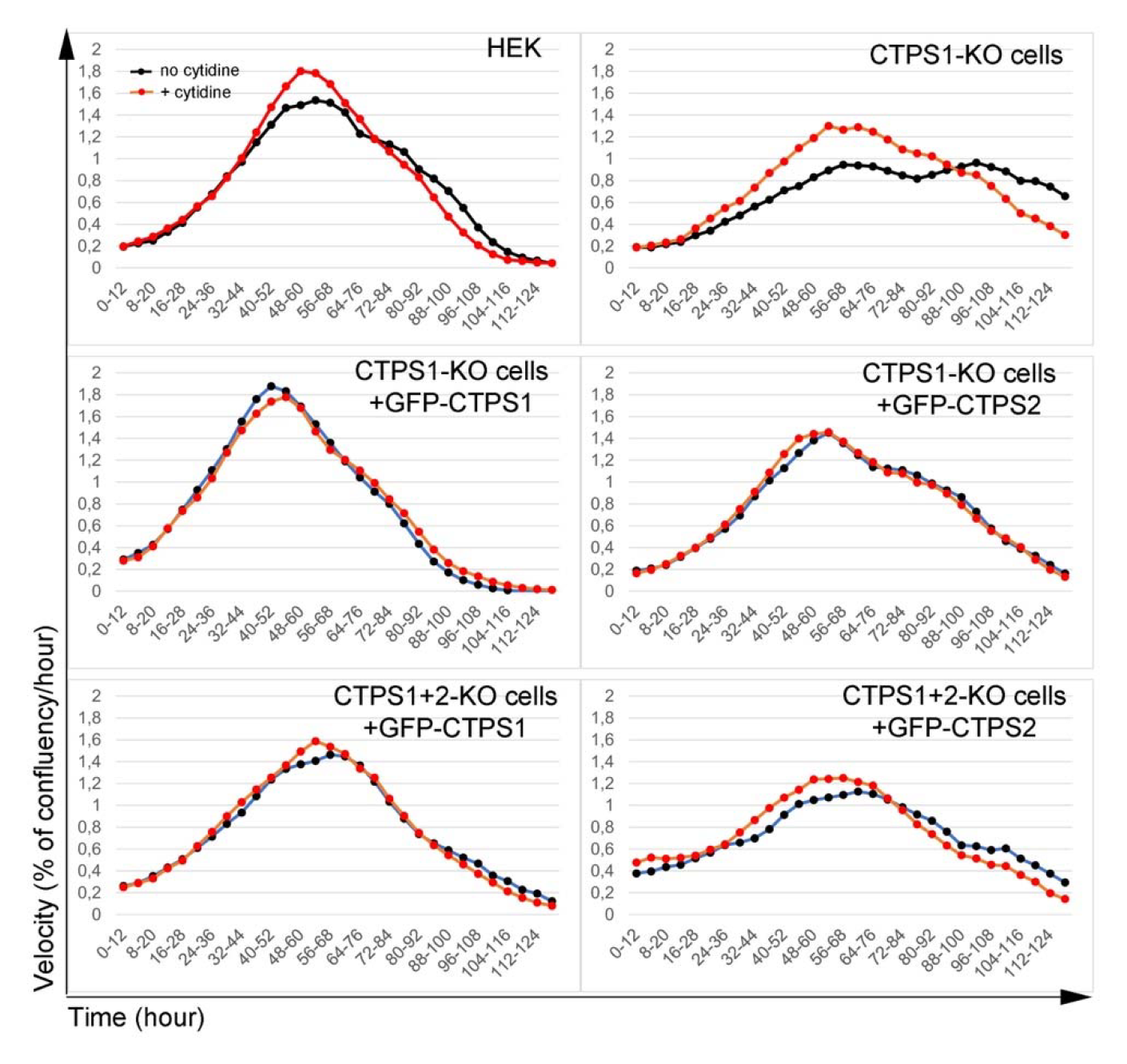
CTPS1 is more efficient than CTPS2 in restoring the proliferation of CTPS1-KO and CTPS1+2-KO HEK cells. Proliferation velocity, corresponding to the Figure 5C. Velocity for each population in presence (red dotted line) or absence (black dotted line) of cytidine supplementation (200µM), corresponding to the increase in confluency per hour on 12-hour periods. Data of one representative experiment of 4 independent experiments except for CTPS1+2KO cells with GFP-CTPS2 only tested 2 times.

**FIGURE S6.**
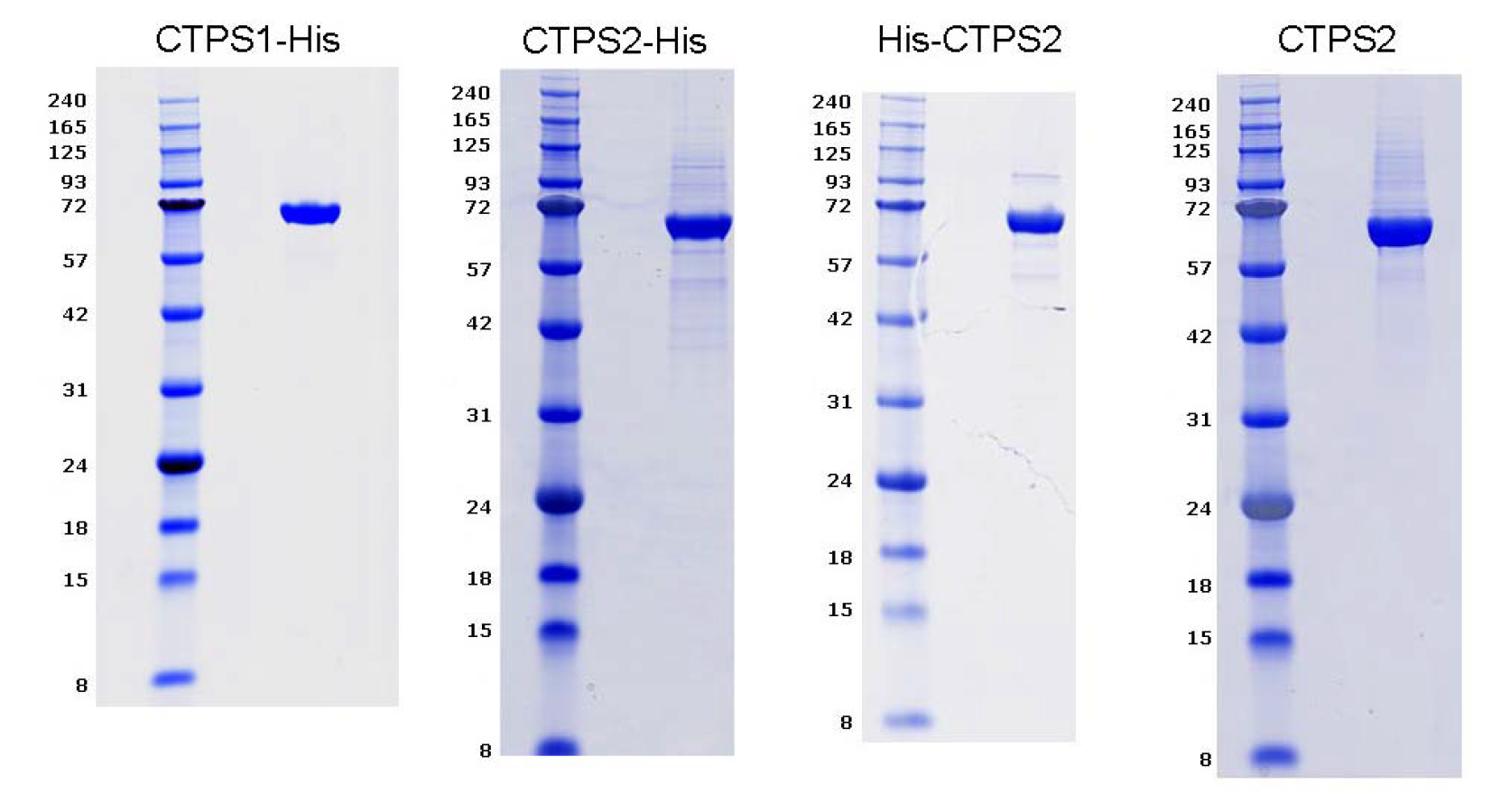
Purified recombinant CTPS1 and CTPS2 proteins. Analysis of purified recombinant tagged forms of CTPS2 and CTPS1 with C-terminal FLAG-Histidine tag (CTPS1- His, CTPS2-His), CTPS2 with N-terminal Histidine tag (His-CTPS2) and CTPS2 without a tag (CTPS2) used in Figure 6 by SDS-PAGE with 10% acrylamide gels. Molecular weight markers on the right. Arrows indicate the purified proteins. The purity is >90% for each protein.

